# High sensitivity quantitative proteomics using accumulated ion monitoring and automated multidimensional nano-flow chromatography

**DOI:** 10.1101/128991

**Authors:** Paolo Cifani, Alex Kentsis

## Abstract

Quantitative proteomics using high-resolution and accuracy mass spectrometry promises to transform our understanding of biological systems and disease. Recent development of parallel reaction monitoring (PRM) using hybrid instruments substantially improved the specificity of targeted mass spectrometry. Combined with high-efficiency ion trapping, this approach also provided significant improvements in sensitivity. Here, we investigated the effects of ion isolation and accumulation on the sensitivity and quantitative accuracy of targeted proteomics using the recently developed hybrid quadrupole-Orbitrap-linear ion trap mass spectrometer. We leveraged ultra-high efficiency nano-electrospray ionization under optimized conditions to achieve yoctomolar sensitivity with more than seven orders of linear quantitative accuracy. To enable sensitive and specific targeted mass spectrometry, we implemented an automated, scalable two-dimensional (2D) ion exchange-reversed phase nano-scale chromatography system. We found that 2D chromatography improved the sensitivity and accuracy of both PRM and an intact precursor scanning mass spectrometry method, termed accumulated ion monitoring (AIM), by more than 100-fold. Combined with automated 2D nano-scale chromatography, AIM achieved sub-attomolar limits of detection of endogenous proteins in complex biological proteomes. This allowed quantitation of absolute abundance of the human transcription factor MEF2C at approximately 100 molecules/cell, and determination of its phosphorylation stoichiometry from as little as 1 μg of extracts isolated from 10,000 human cells. The combination of automated multidimensional nano-scale chromatography and targeted mass spectrometry should enable ultra-sensitive high-accuracy quantitative proteomics of complex biological systems and diseases.

## INTRODUCTION

The emerging ability to measure cellular and physiological states accurately and quantitatively promises to transform our understanding of biology and disease [1]. For example, time-resolved and multi-parametric quantitative analyses of cellular signaling are enabling the elucidation of fundamental paradigms of cell development and homeostasis [2], [3]. Likewise, accurate measurements of human disease states are enabling improved diagnostic markers and refined mechanisms of disease pathophysiology [4], [5].

In large part, these advances were made possible by the development of increasingly accurate and sensitive methods for quantitative analysis of proteins and their post-translational modifications in complex biological proteomes. For example, selected reaction monitoring (SRM) utilizes quadrupole mass analyzers to filter specific precursor and fragment ions produced by collision-induced dissociation (CID) [6]-[9]. This method benefits from high-efficiency continuous ion beams, and relatively high sensitivity of direct dynode detection, but is subject to interference effects due to the relatively low unit-mass resolution of quadrupole mass analyzers. As a result, SRM methods require specialized approaches to control for variable specificity, hindering their widespread use [7], [10]-[12].

To overcome these limitations, parallel reaction monitoring (PRM) has been developed by leveraging high-resolution Orbitrap mass analyzers to improve assay specificity as a result of monitoring fragment ions with parts per million (ppm) mass accuracy [13], [14]. Likewise, the incorporation of high-resolution time-of-flight (TOF) mass analyzers has been used to improve the accuracy of reaction monitoring methods, including their use in data-independent approaches such as SWATH [15], [16]. Thus far, targeted mass spectrometry methods exhibit at least 10-fold better sensitivity than data-independent approaches [15]. Consequently, recent efforts have focused on improving the ion transfer efficiencies of these methods, such as the recently introduced parallel accumulation-serial fragmentation technique [17].

The requirement for high ion transfer efficiencies for accurate quantitative proteomics led to the incorporation of ion trapping devices in modern mass spectrometry instruments. For example, the use of external ion storage improved the accumulation efficiency of Fourier transform-ion cyclotron resonance (FT-ICR) mass spectrometers [18], [19]. Likewise, implementation of Orbitrap mass analyzers necessitated the incorporation of external ion storage devices with improved electrodynamic concentration properties [20], as originally required for coupling bright continuous ion sources to ICR mass analyzers [21]. Recent implementations of ion storage on hybrid instruments, such as the Q Exactive quadrupole-Orbitrap (Q-OT), and the Fusion quadrupole-Orbitrap-linear ion trap (Q-OT-IT) mass spectrometers, utilize high-capacity multipole ion traps, which permit accumulation and routing of ions prior to analysis [22], [23].

Here, we investigated the effects of ion selection and accumulation on the sensitivity and quantitative accuracy of targeted proteomics using the recently developed hybrid Fusion quadrupole-Orbitrap-linear ion trap mass spectrometer. We leveraged ultra-high efficiency low μm-scale nano-electrospray ionization to determine the absolute limits of sensitivity, achieving yoctomolar absolute limits of quantitation with more than seven orders of linear quantitative accuracy. We observed that ion co-isolation led to significant reduction in sensitivity in analyses of complex human cellular proteomes. To partially overcome this limitation, we implemented an automated, scalable two-dimensional ion exchange-reversed phase nano-scale chromatography system, suitable for robust, high-resolution, high-capacity separations necessary for quantitative targeted mass spectrometry. By quantifying the endogenous transcription factor MEF2C and its phosphorylation stoichiometry in 1 μg of extracts from as few as 10,000 human cells, we achieved significant improvements in the sensitivity and quantitative accuracy of both PRM and accumulation monitoring (AIM) methods, permitting the detection and quantitation of approximately 100 molecules/cell.

## EXPERIMENTAL PROCEDURES

### Reagents

Mass spectrometry grade (Optima LC/MS) water, acetonitrile (ACN), and methanol were from Fisher Scientific (Fair Lawn, NJ). Formic acid of 99%+ purity (FA) was obtained from Thermo Scientific. Ammonium formate and all other reagents at MS-grade purities were obtained from Sigma-Aldrich (St. Louis, MO).

### Synthetic peptides and proteome preparation

MRFA peptide was obtained from Sigma-Aldrich. Based on consensus protein sequences in the UniProt database (as of January 30^th^, 2015), N-terminally isotopically labeled (^13^C_6_^15^N_2_ lysine and ^13^C_6_^15^N_4_ arginine) MEF2C and MARK4 peptides (Table 1) were synthesized using solid phase chemistry by New England Peptides (Gardner, MA), and purified by reversed phase chromatography. Extinction coefficients were calculated as described by Kuipers and Gruppen [24], and listed in Supplementary Table 1. Peptides were quantified using UV absorbance spectroscopy at 214 nm using 3 mm QS quartz cuvettes (Hellma, Plainview, NY) and the SpectraMax M5 analytical spectrophotometer (Molecular Devices, Sunnyvale, CA). A standard peptide mixture was created by mixing individual peptides at final concentration 10 pmol/μl in a clean glass vial. For direct infusion experiments, the peptide mixture was serially diluted 1:10 in 30% ACN, 0.1% FA containing 1 μg/ml MRFA peptide (Sigma-Aldrich).

**Table 1.** **Peptide properties.**

| ID | Sequence | m/z |
| --- | --- | --- |
| M1 | SEPVSPPR | 439.7339 |
| M2 | SEPV(pS)PPR | 479.7171 |
| M3 | NSPGLLVSPGNLNK | 709.3981 |
| M4 | N(pS)PGLLVSPGNLNK | 749.3813 |
| M5 | YTEYNEPHESR | 717.8116 |
| M6 | LFEVIETEK | 558.3073 |
| M9 | VLIPPGSK | 409.7649 |

Human OCI-AML2 cells were obtained from the German Collection of Microorganisms and Cell Cultures (Brunswick, Germany). Cells were cultured as described [25], collected while in exponential growth phase, washed twice in ice-cold PBS, snap frozen and stored at −80^°^C. Protein extraction and proteolysis was performed as previously described [26]. Briefly, frozen pellets of 5 million cells were thawed on ice, resuspended in 100 μl of 6 M guanidinium hydrochloride, 100 mM ammonium bicarbonate at pH 7.6 (ABC) containing PhoStop phosphatase inhibitors (Roche Diagnostics GbmH, Mannheim, Germany), and lysed using the E210 adaptive focused sonicator (Covaris, Woburn, CA). The protein content in cell lysate was determined using the BCA assay, according to the manufacturer’s instructions (Pierce, Rockford, IL). Upon reduction and alkylation, proteomes were digested using 1:100 w/w (protease:proteome) LysC endopeptidase (Wako Chemical, Richmond, VA) and 1:50 w/w MS sequencing-grade modified trypsin (Promega, Madison WI). Digestion was stopped by acidifying the reactions to pH 3 using formic acid (Thermo Scientific), and peptides were subsequently desalted using solid phase extraction using C18 Macro Spin columns (Nest Group, Southborough, MA). Peptides were eluted in 60% acetonitrile, 1% formate in water, lyophilized using vacuum centrifugation, and stored at −20°C. Tryptic peptides were reconstituted in 0.1% formate, 3% acetonitrile to a final concentration of 0.5 μg/μl. For experiments in cellular proteomes, the synthetic peptide mixture was initially diluted to 5 pmol/μl in a tryptic digest of whole OCI-AML2 cell proteome at 0.5 μg/μl, and subsequently serially diluted 1:10 in the same solution.

### Nanoscale liquid chromatography

Detailed description of the instrumental and operational parameters, as well as step-by-step protocol for system construction and operation are provided in Supplementary Materials. Both direct infusion sample delivery and liquid chromatography experiments were performed using the Ekspert NanoLC 425 chromatograph (Eksigent, Redwood city, CA), equipped with an autosampler module, two 10-port and one 6-port rotary valves, and one isocratic and two binary pumps. Polyimide-coated fused silica capillaries (365 μm outer diameter, variable inner diameters) were obtained from Polymicro Technologies (Phoenix, AZ.) Unions and fittings were obtained from Valco (Houston, TX). For direct infusion, samples were initially aspirated into a 10 μl PEEK sample loop. Upon valve switching, the content of the loop was ejected using a gradient pump (30% ACN, 0.1% FA at 100 nl/min) into an empty silica capillary (20 μm inner diameter) in-line with the DPV-566 PicoView nano-electrospray ion source (New Objective, Woburn, MA).

Chromatographic columns were fabricated by pressure filling the stationary phase into silica capillaries fritted with K-silicate, as previously described [27]. Strong-cation exchange columns were fabricated by packing Polysulfoethyl A 5 μm silica particles (PolyLC, Columbia, MD) into 150 μm × 10 cm fritted capillary. Reversed phase columns were fabricated by packing Reprosil 1.9 μm silica C18 particles (Dr. Meisch, Ammerbauch-Entrigen, Germany) into 75 μm × 40 cm fritted capillaries. Trap columns were fabricated by packing Poros R2 10 μm C18 particles (Life Technologies, Norwalk, CT) into 150 μm × 4 cm fritted capillaries.

Vented trap-elute architecture was used for chromatography [28]. One of the 10-port valves was set to include in the flow path either the SCX column or an empty capillary of equal inner volume. Samples were initially loaded into a 10 μl PEEK sample loop, and subsequently delivered at 1 μl/min by the isocratic pump using 0.1% FA in water into either the empty capillary (for one-dimensional chromatography) or the SCX column (for two-dimensional chromatography). A step gradient of 50, 100, 150, 300, and 1000 mM ammonium formate (AF) in water, pH 3 was delivered in 3.5 μl (0.5 column volume) increments from auto-sampler vials to elute peptides into the trap column, where peptides were desalted. Lastly, peptides were resolved by reversed phase chromatography hyphenated to the nano-electrospray ion source. Upon valve switch to connect the trap column in line with the analytical reversed phase column and ion emitter, the pressure was equilibrated at a flow of 250 nl/min for 5 minutes in 5% buffer B (ACN, 0.1% FA) in buffer A (water, 0.1% FA). Subsequently, a 60-minutes linear gradient of 5-38% of buffer B was used to resolve peptides, followed by a 5 minutes 38-80% gradient prior to column wash at 80% buffer B for 30 minutes.

### Electrospray ionization and mass spectrometry

Electrospray emitters with terminal opening diameter of 2-3 μm were fabricated from silica capillaries as previously described [29]. The emitter was connected to the outlet of the reversed phase column using a metal union that also served as the electrospray current electrode. Electrospray ionization was achieved using variable voltage, programmed from 1750 to 1450 V with 50 V steps over 60 minutes of the gradient elution. During column loading, the electrospray emitter was washed with 50% aqueous methanol using the DPV-565 PicoView ion source (New Objective).

For all measurements, we used the Orbitrap Fusion mass spectrometer (Thermo Scientific, San Jose, CA). During AIM measurements, the mass spectrometer was programmed to iteratively perform precursor ion scans with 8 Th isolation windows targeting both endogenous light and synthetic heavy peptides using Q1 isolation and S-lens voltage of 60 V [22]. Unless otherwise specified, ions were accumulated for a maximum of 200 ms with automatic gain control of 10^5^ ions, and scanned at 240,000 resolution. For PRM scans, precursor ions were isolated using 2 Th isolation windows, and fragmented by HCD with normalized collision energy set at 32% before analysis of the fragment ions in the Orbitrap at 30,000 resolution. Optimal fragmentation conditions were preliminarily established for each target peptide by manual inspection of MS2 spectra collected within the same analysis upon fragmentation with HCD energy 30, 32, 36, and 38% [30].

### Data Analysis

Ion signal intensities and total ion currents were analyzed using XCalibur Qual Browser 3.0 (Thermo Fisher Scientific). Automated chromatographic peak area integration was performed using Skyline 3.5.0 [31], with mass tolerance set at 0.0075 Da corresponding to 10 ppm for m/z of 750 Da, and integration boundaries were manually verified for all peaks. Numerical and statistical analyses were performed using Origin Pro 9.0 (OriginLab Corporation, Northampton, MA). All raw and processed mass spectrometry data as well as Skyline chromatogram documents are available via ProteomeXchange [32] with identifier PXD006236. Quantitative data from serially diluted synthetic peptides were linearly fitted to obtain a signal-response function for each peptide. These functions were subsequently used to calculate amounts of endogenous peptides. Phosphorylation stoichiometry was defined as the fraction of each peptide being chemically modified, as described [33]. The protein content of OCI-AML2 cells was established by BCA assay quantification of total protein extracted from cells, which were manually counted using a Neubauer hemocytometer.

### Experimental design and statistical rationale

The study evaluated the sensitivity and specificity of targeted detection using peptides delivered either by direct infusion or by chromatography with variable peak capacities. Technical variability was established under direct infusion regime by collecting seven measurements for each data point. For chromatographically resolved peptides, triplicate measurements of the intensity of synthetic peptides were performed at three experimental conditions. Endogenous peptide measurements were performed in parallel with isotopologue targeting, for a total of 18 replicate measurements. To compare AIM and PRM, assays were performed within the same experiment to control for possible variation in chromatography and ionization performance.

## RESULTS

### Accumulated Ion Monitoring for Targeted Proteomics using Hybrid Quadrupole-Orbitrap-Linear Ion Trap (Q-OT-IT) Instrument

Sensitivity of mass spectrometric detection is in principle determined by two factors: the minimum number of ions necessary to produce a measureable electronic signal, and the baseline instrumental noise. In the case of high-resolution mass analyzers coupled to external ion storage devices, the number of ions delivered to the mass analyzer can thus be increased by prolonged ion accumulation prior to detection. The incorporation of high-capacity ion routing multipoles (IRM) on the recently developed Q Exactive (Q-OT) and Fusion (Q-OT-IT) mass spectrometers permits ion accumulation and storage for as long as 5 seconds.

An acquisition strategy was thus designed, combining quadrupole precursor ion filtering, and precursor and fragment ion detection in the Orbitrap and linear ion trap, respectively. This method combines high sensitivity and specificity for targeted mass spectrometry. Such a method is conceptually related to quadrupole-filtered selected ion monitoring (SIM) [34], but because of its use of IRM ion storage and accumulation, we term it accumulated ion monitoring (AIM). AIM is also related to parallel reaction monitoring (PRM) [13], [14], insofar as both methods achieve high-specificity quantification through high-resolution detection of trapped ions, but is distinguished by the identification of ions using parallel fragment ion scanning in the linear ion trap, as specifically enabled by the so-called tribrid architecture of the Fusion Q-OT-IT instrument [22], [23].

### Establishing the Limits of Quantitation and Quantitative Accuracy of AIM

In principle, extended ion accumulation of AIM and PRM could enhance the detectability of peptides and extend the quantitative accuracy and overall sensitivity of targeted proteomics. To test this hypothesis, we determined the absolute sensitivity of AIM detection, and its dependence upon ion accumulation and storage, using synthetic peptides delivered by continuous infusion. We initially chose tryptic peptides derived from human transcription factor MEF2C and kinase MARK4 given their physicochemical and ionization properties that are generally representative of human tryptic peptides (Table 1, Supplementary Table 1). To control for possible adsorptive losses, peptides were serially diluted in neat solvent containing the MRFA peptide as a carrier. To maximize nano-electrospray ionization efficiencies while maintaining robust emitter performance [35], [36], we fabricated fused silica emitter tips with tip openings of 2-3 μm, as determined using scanning electron and optical microscopy (Supplementary Figure 1A & B) [29]. This enabled ionization efficiencies of 3-23% for the monitored peptides, as estimated by the time required to achieve the automatic gain control (AGC) threshold values (Supplementary Figure 2).

Using these parameters, we analyzed the absolute sensitivity of AIM by measuring the signal response of synthetic peptides serially diluted in MRFA-containing neat solvent. Remarkably, this achieved specific detection of less than 1 ymol/ms of peptides, corresponding to less than 100 ions per scan, with nearly 7 orders of magnitude of linear quantitative accuracy (Figure 1A, Supplementary Figures 3, Supplementary Table 2). We observed no significant differences in the limits of quantitation by increasing the ion accumulation time from 25 to 2500 ms (Figure 1A).

**Figure 1.**
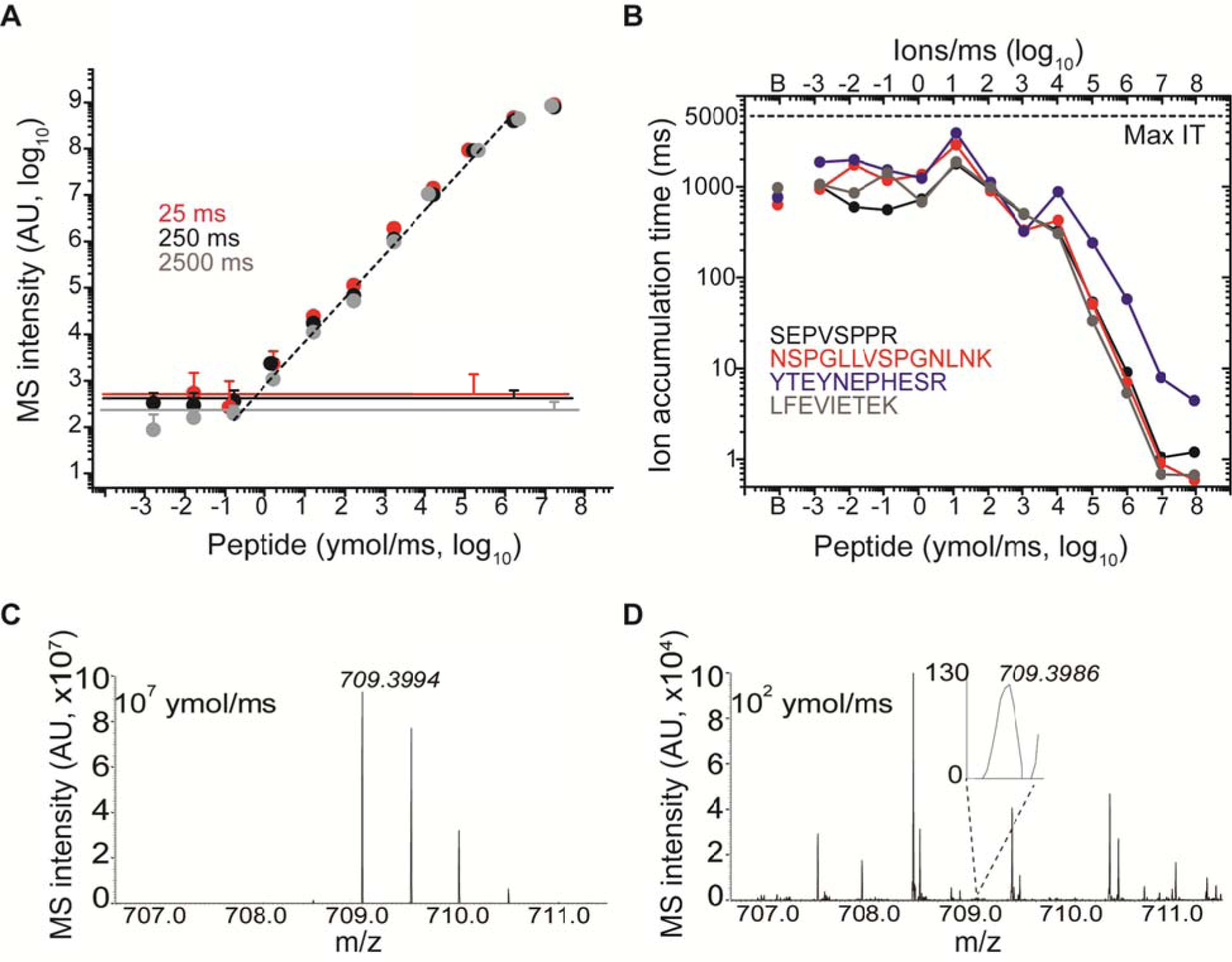
A) AIM acquisition enables detection of 2-5 yoctomoles/ms of target peptides serially diluted in neat solvent, and seven order of magnitude of linear dynamic range. The MS intensity within 10 ppm from the theoretical m/z was recorded for peptide NSPGLLVSPGNLNK (m/z 709.3981) with maximum injection time (max IT) 25 (red), 250 (black), and 2500 (grey) ms (n=7, error bars represent standard deviation of independent experiments). Noise levels recorded at baseline level (i.e., with no target peptide infused) is displayed for each maximum injection times as continuous line. Overlapping data points were horizontally offset when needed for clarity. B) Ion accumulation times obtained for each target peptide with maximum injection time set at 5s. Ion injection time in the baseline samples (i.e. with no target present) is denoted as “B”. Comparison of spectra recorded with target flow 10e7 (C) and 100 ymol/ms (D) reveals that target accumulation is practically limited by co-isolated contaminant ions (ion intensity on absolute linear scale).

Analysis of ion accumulation times under non-limiting maximum injection times revealed two distinct regimes of operation (Figure 1B). As long the target peptide was the dominant ion in the isolation window, i.e. delivered at flow rates greater than approximately 10 zmol/ms, ion accumulation times scaled linearly with peptide concentration (Figure 1B). However, for peptide targets delivered at less than 10 zmol/ms, contaminant ions from solvent, silica and ambient air limited accumulation times to approximately 2 seconds (Figure 1B), as confirmed by spectral analysis (Figure 1C & D). Thus, AIM can achieve sensitivities on the order of 100 ions/scan, but target co-isolation substantially limits target ion accumulation, even under optimized analytical conditions.

### AIM Quantitative Proteomics using Automated Two-Dimensional Nanoscale Chromatography

Since peptide target purity affects ions accumulation, which would be further exacerbated in the analysis of complex biological proteomes, we sought to enhance the sensitivity of ion trap-targeted method by reducing contaminant co-isolation. Co-isolation is normally controlled by gas phase fractionation using quadrupolar mass filters [37]. However, this strategy is potentially affected by variable transmittance, particularly for narrow selected m/z ranges, and may introduce biases from independent selection of isotopologues when the isolation window is smaller than their m/z difference. In addition, we reasoned that high-resolution chromatography [38] could be used to improve the isolation of specific peptides and consequently their sensitive quantitation by both AIM and other targeted proteomics methods such as PRM. Based on prior implementations of multi-dimensional separations [28], [39]-[43] we designed an automated two-dimensional (2D) strong cation exchange (SCX) and reversed phase (RP) nano-scale chromatography architecture (Figure 2). This implementation benefits from scalability, allowing for seamless integration of additional separation dimensions, and direct interoperability between multi-dimensional and one-dimensional (1D) modes of operation, allowing for rigorous analysis of mass spectrometric quantitative performance (Figure 3). Reduction of SCX chromatographic resolution was previously described as a result of isocratic mass elution, particularly during step gradient elution [44]. To address this issue we devised an optimized method that minimizes the volumes of mobile phase flowing through the SCX column (Supplementary Methods). Detailed description of the instrumental features, as well as step-by-step construction and operation instructions are provided in the Supplementary Materials.

**Figure 2.**
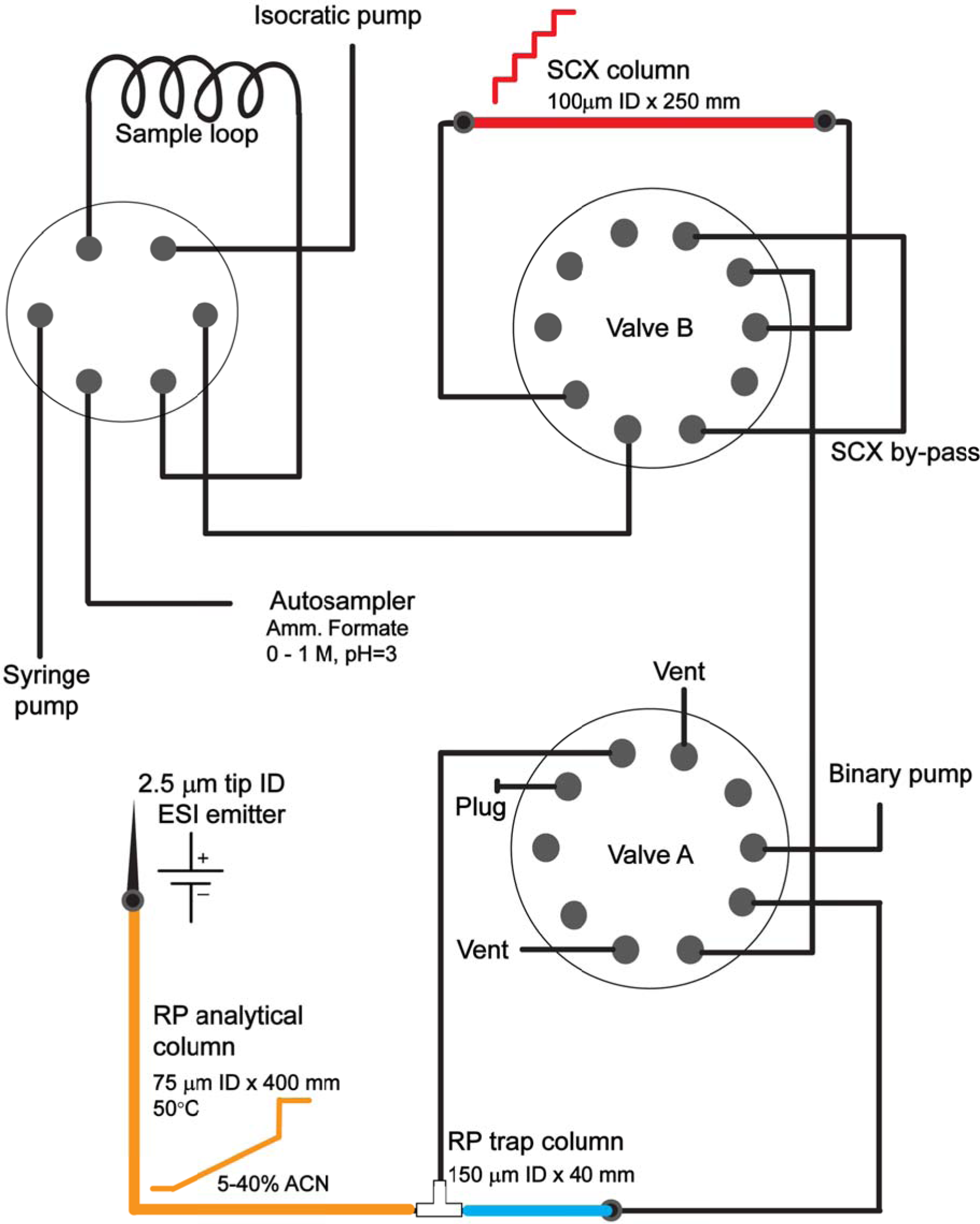
Overview of the automated 2D chromatography system. The system involves #describe the key components succinctly

**Figure 3.**
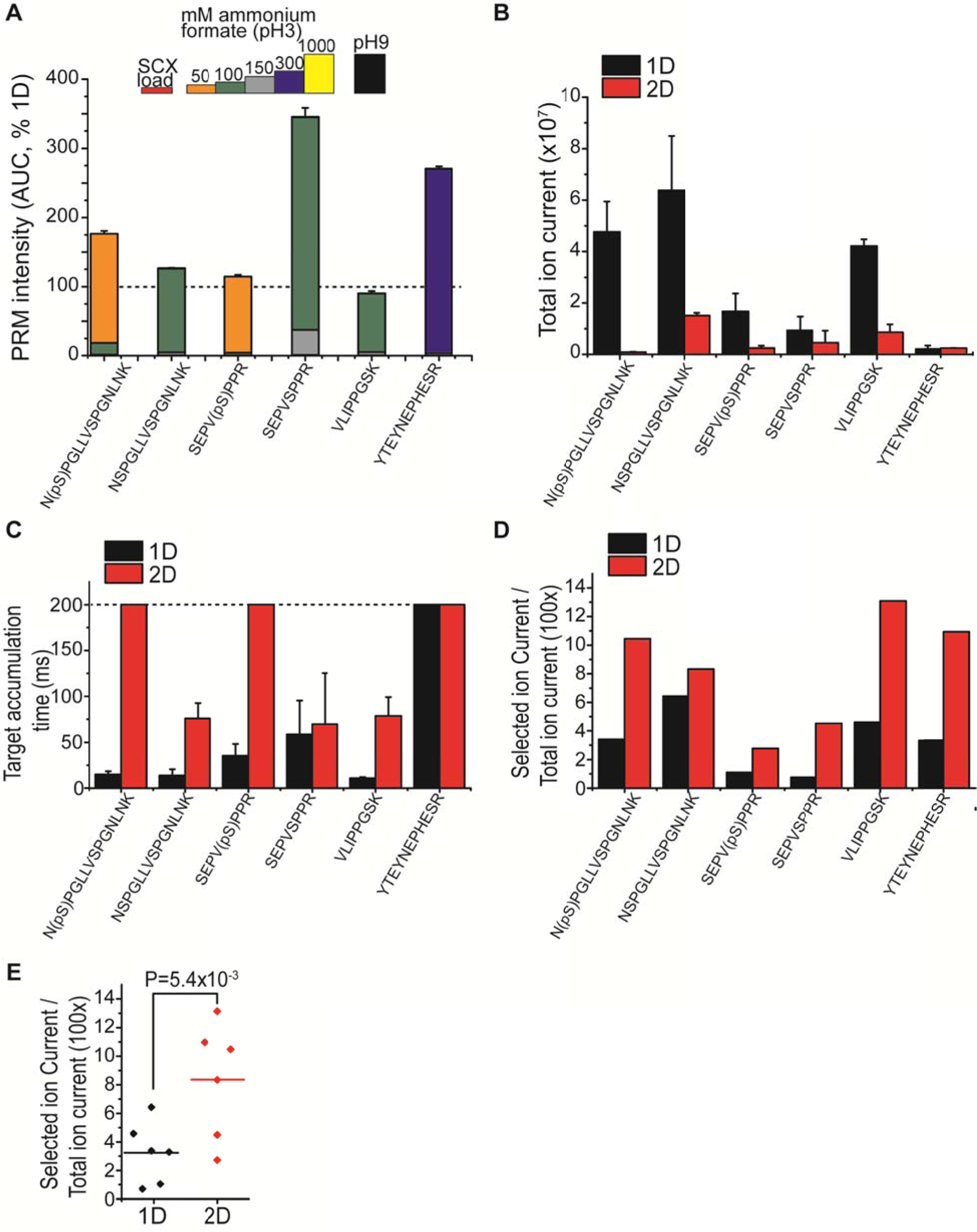
A) Automated 2D chromatography system enables efficient resolution of target peptides. Average recovery and SCX fractionation efficiency of peptides are plotted relative to 1D (n=4, error bars represent standard deviation of fraction of target in best SCX fraction. Quantification by PRM). B) 2D chromatography reduces total ion current from co-isolated contaminant ions, in absence of synthetic targets (n=3, error bars = standard deviation). C) 2D chromatography improves ion accumulation, in absence of synthetic targets (n=3, error bars = standard deviation). Maximum ion injection time (200 ms, dashed line) is achieved for phosphorylated peptides. D) 2D chromatography improves the ratio of MS signal specific for target ions (specific ion current), over total ion current within the scan (10 fmol target peptide, currents at apex of chromatographic peak). E) Improvement of SIC/TIC by SCX-RP chromatography is statistically significant (paired t-test P=0.0054, n=6).

To establish the chromatographic performance of this system, we analyzed the separation efficiency of synthetic isotopically-labeled targets diluted in 1 μg of tryptic peptides isolated from human cells, analyzed under identical conditions and operational parameters in 2D as compared to 1D separations. First, we assessed the recovery of separated peptides by comparing their SCX retention as fractionated by step-wise elution with increasing concentrations of ammonium formate at pH 3, with the complete column elution from 1D chromatography (Figure 3A). Using this approach, we observed essentially complete recovery of target peptides in 2D versus 1D separations, with minimal losses due to incomplete retention, as assessed by analysis of the flow-through fractions (Figure 3A). Notably, the two most hydrophilic peptides exhibited significantly increased (2.6-3-fold) signal intensities in 2D as compared to 1D separations, consistent with their improved retention by the final reverse phase column upon SCX fractionation (Figure 3B). We found that retention times in the final reverse phase separations were highly reproducible for 2D-separated peptides, with a constant delay of 30-90 seconds, as compared to 1D-separated analytes (Supplementary Figure 4). Peptide LFEVIETEK was not recovered from the reverse phase column regardless of pre-fractionation due to poor retention properties, and was therefore omitted in further analyses. Importantly, 2D separation enabled nearly complete (93% average) resolution of target peptides in individual SCX fractions, as assessed by the analysis of signal intensities detected across fractions (Figure 3A). Using recent formalizations of multi-dimensional chromatographic separations [45], we estimated that our current 2D implementation produced approximately 6-fold increase in the chromatographic efficiency, as compared to conventional 1D chromatography alone.

To test the hypothesis that automated 2D nano-scale chromatography can reduce co-elution of background ion and thus their co-isolation in AIM and PRM, we compared the total ion currents (TIC) observed at the expected retention times of target peptides in 1 μg of tryptic human cell extract, analyzed in 2D as compared to 1D but otherwise under identical conditions. Consistent with the expected reduction of contaminant co-isolation, we observed significant reduction of TIC levels for most target peptides (Figure 3B). We found that target ion accumulation, as assessed by the effective injection time, significantly increased for most target peptides under 2D as compared to 1D separation (Figure 3C). Lastly, we estimated the effective purification of specific target peptides by analyzing their fractional specific ion current (SIC) as compared to the total ion current (TIC). This analysis showed that the SIC/TIC ratios for most target peptides were significantly increased by 2D as compared to 1D separations (average > 3-fold, *p* = 5.4e-3, t-test, Figures 3D & E).

### Automated Two-Dimensional Nano-scale Chromatography Improves Sensitivity and Quantitative Accuracy of AIM and PRM Targeted Proteomics

Having confirmed improved target accumulation by the automated 2D chromatography, we next sought to determine its effect on the sensitivity of targeted detection in complex biological proteomes. To test this, we used AIM and PRM to quantify synthetic isotopically-labeled peptides (Table 1) serially diluted in 1 μg of tryptic human cell proteome (Figure 4A & B, Supplementary Figure 5). For AIM, we used a quadrupole isolation window of 8 Th to enable simultaneous and unbiased co-isolation of both endogenous (light) and synthetic (heavy) peptides, and compared the performance of automated 2D versus 1D separations under otherwise identical conditions. Targeted fragment spectra under optimized HCD condition were recorded for PRM along with AIM scanning within the same experiment, thus enabling unbiased comparison of the two methods.

**Figure 4.**
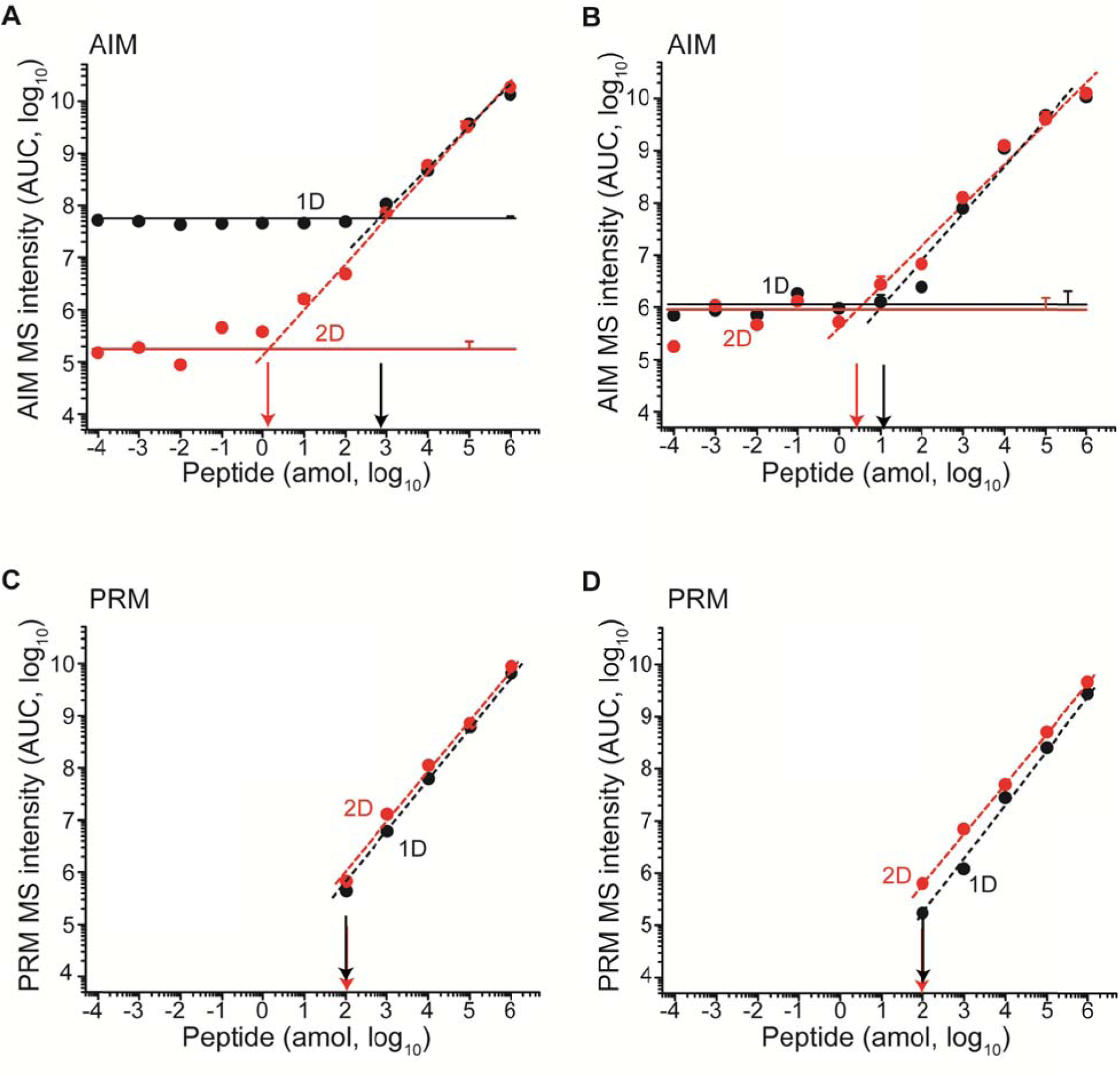
The improvement in sensitivity from 2D depends on efficiency of fractionation. Phosphorylated peptide N(pS)PGLLVSPGNLNK is efficiently resolved from isobaric contaminants, resulting in a 2 amol AIM sensitivity after SCX compared to 1 fmol in 1D. PRM analysis achieves 100 amol LOD both in 1D and 2D. Peptide NSPGLLVSPGNLNK is poorly resolved by SCX from the bulk of doubly charged tryptic peptides, resulting in similar sensitivity in 1D (black) and 2D (red) by both AIM (C) and PRM (D). Horizontal lines represent average noise levels (n=3) for 1D (black) and 2D (red) in AIM. Error bars represent technical variability as standard deviation (n=3) measured at noise level, 10 amol and 100 fmol. Overlapping data points are horizontally offset when needed for clarity. Vertical arrows mark the LOD in 1D (black) and 2D (red) respectively.

We observed that for some peptides, and particularly phospho-peptides which are efficiently resolved by SCX chromatography, AIM 2D chromatography achieved sub-attomolar limits of detection, with an improvement of nearly 3 orders of magnitude as compared to 1D (Figure 4A, Supplementary Figure 5C). However, this improvement was not universal, as doubly charged peptides that tend to co-elute with the majority of human tryptic peptides, exhibited similar limits of detection (Figure 4B, Supplementary Figure 5), in agreement with their similar fractional SIC/TIC values in 1D and 2D (Figure 3D). Nonetheless, even for these peptides, AIM achieved limits of detection in the 1-10 attomolar range (Figure 4B, Supplementary Figure 5). In agreement with prior studies [14], PRM achieved limits of detection in the 10-100 attomolar range (Figure 4C & D), consistent with its dependency on ion fragmentation for quantitation (defined as at least 3 fragment ions in 5 consecutive scans), in contrast to AIM which only uses fragment ions for identification. Similarly, we found that automated 2D chromatography improved the sensitivity of PRM detection of some peptides to the 10 attomol range (Supplementary Figure 5). In all, automated 2D chromatography improved the sensitivity and quantitative accuracy of AIM and PRM by nearly 100-fold with sub-attomolar sensitivity in complex biological proteomes.

### Quantitative Proteomics with Ultra-High Accuracy and Precision using Automated 2D AIM and PRM

To establish the analytical performance of automated 2D chromatography for quantitative proteomics, we determined the absolute abundance of endogenous peptides derived from the master transcription factor MEF2C [46], including its two functional phosphorylation sites, in human leukemia cells OCI-AML2. We analyzed 1 μg of tryptic peptides extracted from 10,000 cells, having diluted isotopically-labeled synthetic peptides as reference standards for absolute quantitation. We found that both automated 2D AIM and PRM accurately quantified the abundance of non-modified endogenous MEF2C peptides, at the mean level of 180 amol/μg total cell lysate, corresponding to on average 11,000 molecules/cell (Figure 5). Consistent with the improved sensitivity of 2D versus 1D targeted proteomics (Figures 3 & 4), more targeted peptides were accurately quantified by 2D AIM and PRM, as compared to their 1D versions (Figure 5A & B). Likewise, 2D AIM exhibited superior sensitivity as compared to 2D PRM, particularly for phospho-peptides, one of which (MEF2C SPEV(pS^396^)PPR) was not detectable by 2D PRM, due to its low attomolar abundance, corresponding to approximately 100 molecules/cell (Figure 5A & B). Notably, 2D AIM enabled both quantitation of absolute abundance and phosphorylation stoichiometry for endogenous MEF2C S222, achieving accurate mean measurements of 42 amol of phosphorylated peptide per μg total cell lysate, with phosphorylation stoichiometry of 26% (Figures 5A & B) [33]. Finally, whereas AIM exhibited improved sensitivity, PRM had comparatively lower variability, partly because of reduced mass accuracy for highly concentrate and pure target peptides, which reduced accuracy of peak profiling (Supplementary Figure 6-7). Thus, automated two-dimensional nano-scale chromatography enables ultra-sensitive high-accuracy quantitative mass spectrometry of complex biological proteomes.

**Figure 5.**
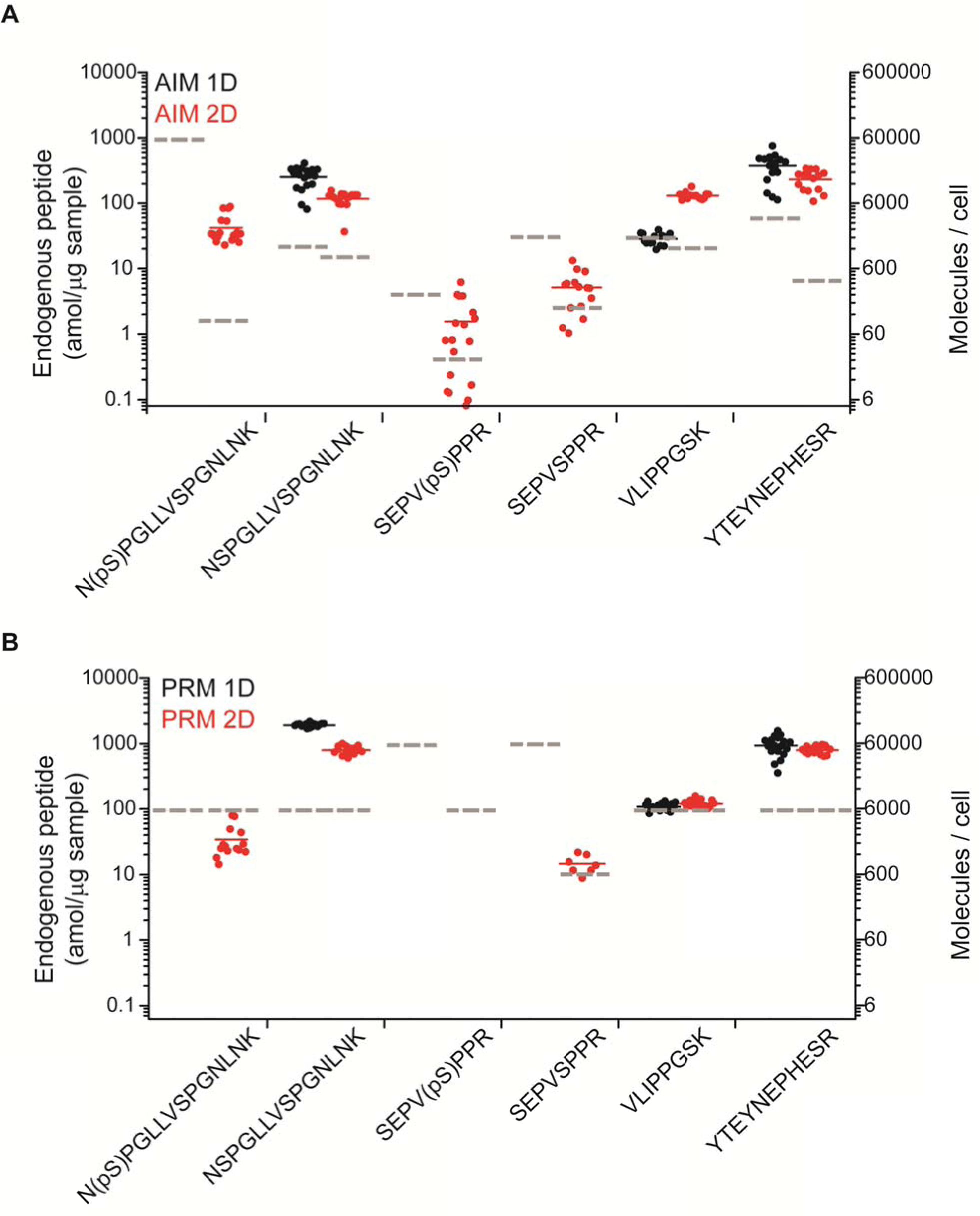
Cellular amount of endogenous peptides from protein MEF2C in 1 μg sample, from approximately 10,000 cells, as determined by 1D (black) and 2D (red) using A) AIM and B) PRM acquisition. Quantification of endogenous phosphorylated peptides was exclusively achievable using SCX. Each dot represents an independent measurements (full horizontal lines indicate average). LOD for each assay, established from serially diluted isotopologues, is denoted with dashed lines in correspondence.

## DISCUSSION

Continuous improvements in chromatography and mass spectrometry instrumentation have now established targeted mass spectrometry as a sensitive and reliable tool to elucidate cellular functions and their regulatory mechanisms [47]. Nonetheless, limits of detection of current implementations of targeted mass spectrometry, combined with the low microgram loading capacity of nano-scale chromatographic columns, imply that quantitative analysis of human proteomes is mostly limited to proteins with medium and high cellular copy number [48]. Even for such targets, the sensitivity of current methods is often insufficient to detect peptides bearing low stoichiometry post-translational modifications (PTMs), which may also deteriorate ionization and fragmentation efficiencies of peptides [49], [50].

Our current work systematically examined factors affecting the sensitivity and quantitative accuracy of targeted mass spectrometry using the recently developed hybrid Fusion quadrupole-Orbitrap-linear ion trap mass spectrometer. First, we found that using ultra-high efficiency electrospray ionization under optimized conditions, current mass analyzer sensitivity and ion transmission enable detection of less than one thousand ions with up to seven orders of magnitude of linear quantitative accuracy. Considering the low-μg loading capacity of current nano-scale chromatographic systems, this sensitivity *per se* would be sufficient to detect molecules present at a few copies per cells or isolated from single cells. However, we also found that co-isolation of background ions nearly isobaric to the target peptides practically impedes extended accumulation prior to detection. In consideration of the excellent transmission efficiency and detector sensitivity of current state-of-the-art mass spectrometers, co-trapping of contaminants is thus the principal factor limiting sensitivity of targeted methods such as AIM and PRM.

Current targeted mass spectrometry methods generally rely on fragment ions generation, as this enables the specificity necessary for the analysis of complex proteomes. Here, we demonstrate that targeted analysis of intact precursor ions, termed accumulated ion monitoring (AIM), enables sensitivity equal or superior to that of optimized parallel reaction monitoring methods, with specificity of detection controlled by isotopically encoded reference standards and parallel fragment ion identification. AIM acquisition enables simultaneous and unbiased isolation of both endogenous targets and reference isotopologues, which can be leveraged to reduce duty cycle requirements as compared to current PRM implementations for rare analytes that may require extended ion accumulation.

To reduced contaminant co-isolation and thus facilitate quantitative proteomics of complex biological proteomes, we implemented an automated two-dimensional nano-scale chromatography architecture, based on prior developments in multi-dimensional separations [28], [38]-[41]. Unlike previous implementation of 2D chromatography that were deployed in combination with data-dependent MS acquisition, operation of our setup was specifically optimized to achieve reproducible and high-resolution sample pre-fractionation suitable for quantitative mass spectrometry. This automated 2D chromatography system, accompanied by detailed construction and operation instructions described in the Supplementary Materials, also permits scalable and seamless chromatography mode switching. Automated 2D chromatography improved the detection limits of targeted analysis of both precursor and fragment ions by effectively reducing co-elution of isobaric contaminants in the final reverse phase chromatography. Such improvement depends on the specific retention properties of each target, and strong cation exchange chromatography was particularly effective in improving, as expected, the analytical exposure of phosphorylated peptides. This enabled detection of endogenously phosphorylated peptides from 1 μg of extracts from as little as 10,000 cells, a scale that is mostly unsuitable for conventional phospho-proteomics methods [51]. This proof-of-concept suggests that targeted quantification of specific classes of peptides could be achieved by leveraging high-resolution, high-capacity chromatographic separations orthogonal to reverse phase under acidic conditions [44], [52], [53], without dedicated affinity enrichment.

In addition, robust reproducibility and high efficiency of fractionation using automated 2D chromatography permit a multidimensional retention time scheduling of targeted assays, with consequent increase in the number of measurable targets per experiment at constant duty-cycle. This increase in assay scheduling capacity further expands the scope of recently described strategies such as isotopologue-triggered scanning [54], and can be coupled to ultra-high capacity chromatography [38] as recently described [55]. As a result the number of targeted assays that can be deployed per injection should increase from the hundreds to the thousands. Furthermore, nano-scale 2D chromatography enables efficient fractionation of low-μg samples, such as those obtained from primary samples and disease specimens, enabling targeted analysis of rare cell populations. Multidimensional fractionation can reduce throughput as compared to singly dimensioned analyses. However, the increased sensitivity and breadth of targeting afforded by automated multi-dimensional separation should permit comprehensive panels of targeted MS assays to elucidate of rare regulatory events that control functional biological processes.

## Author Contributions

P.C. and A.K. conceived and designed the study, P.C. performed the experiments, P.C. and A.K. performed analyses, P.C. and A.K. wrote the manuscript.

We thank Scott Ficarro and Jarrod Marto for technical advice, Michael Senko for critical discussion, Avantika Dhabaria for technical assistance, Christopher Lima and Laurent Cappadocia for spectrophotometric analyses, and Nina Lampen for electron microscopy. This work was supported by the NIH R21 CA188881, R01 CA204396, P30 CA008748, Burroughs Wellcome Fund, Josie Robertson Investigator Program, Rita Allen Foundation, Alex’s Lemonade Stand Foundation, American Society of Hematology, and Gabrielle’s Angel Foundation (A.K.), and American Italian Cancer Foundation (P.C.). A.K. is the Damon Runyon-Richard Lumsden Foundation Clinical Investigator.

This article contains Supplemental Material.

## Competing Financial Interests

The authors have no competing financial interests.

## SUPPLEMENTARY FIGURES LEGENDS

**Supplementary Figure 1.**
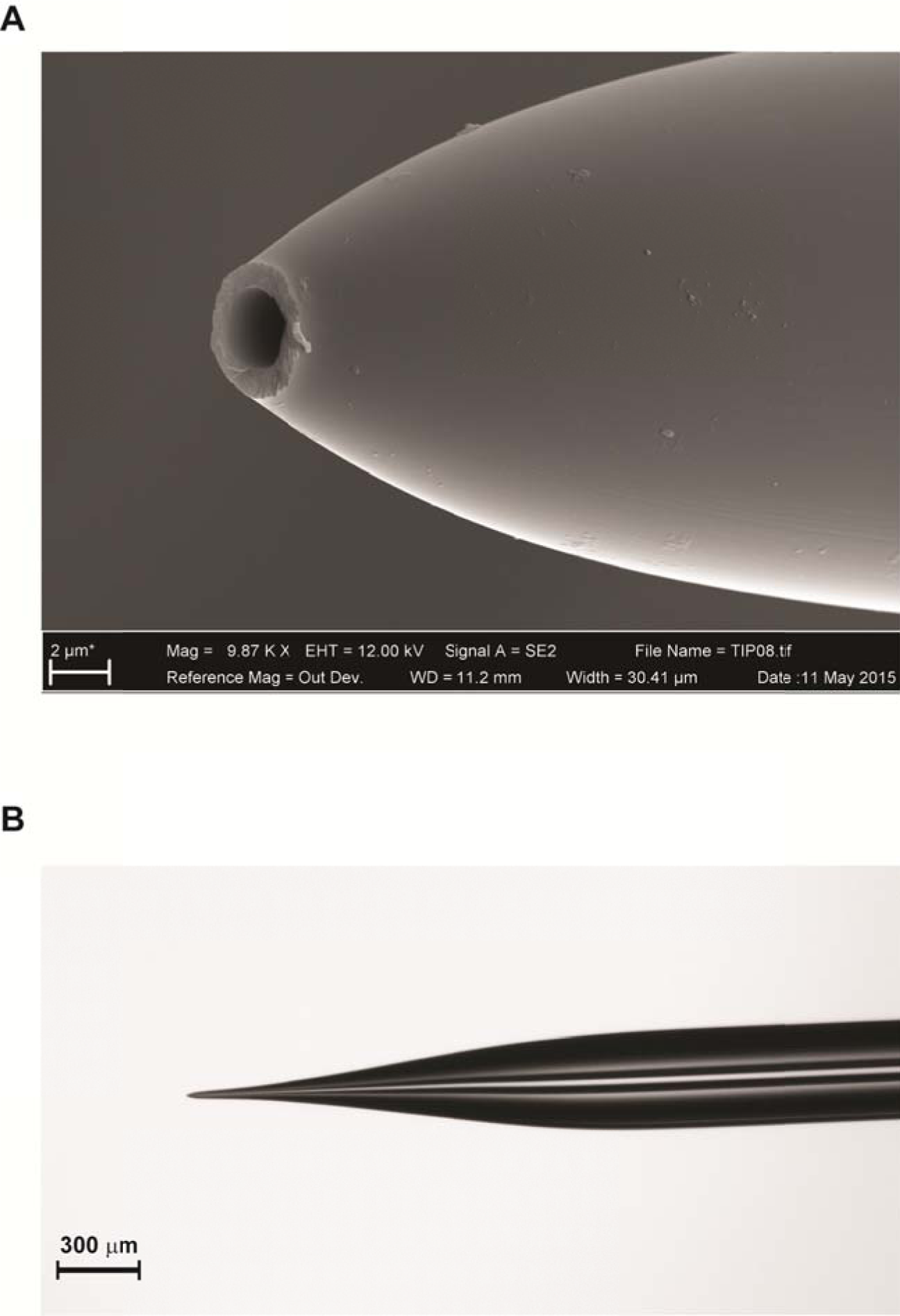
A) Scanning electron micrograph of a typical nano-electrospray emitter tip, with 2 μm tip diameter. B) Photomicrograph (4X) of a typical electrospray tip.

**Supplementary Figure 2.**
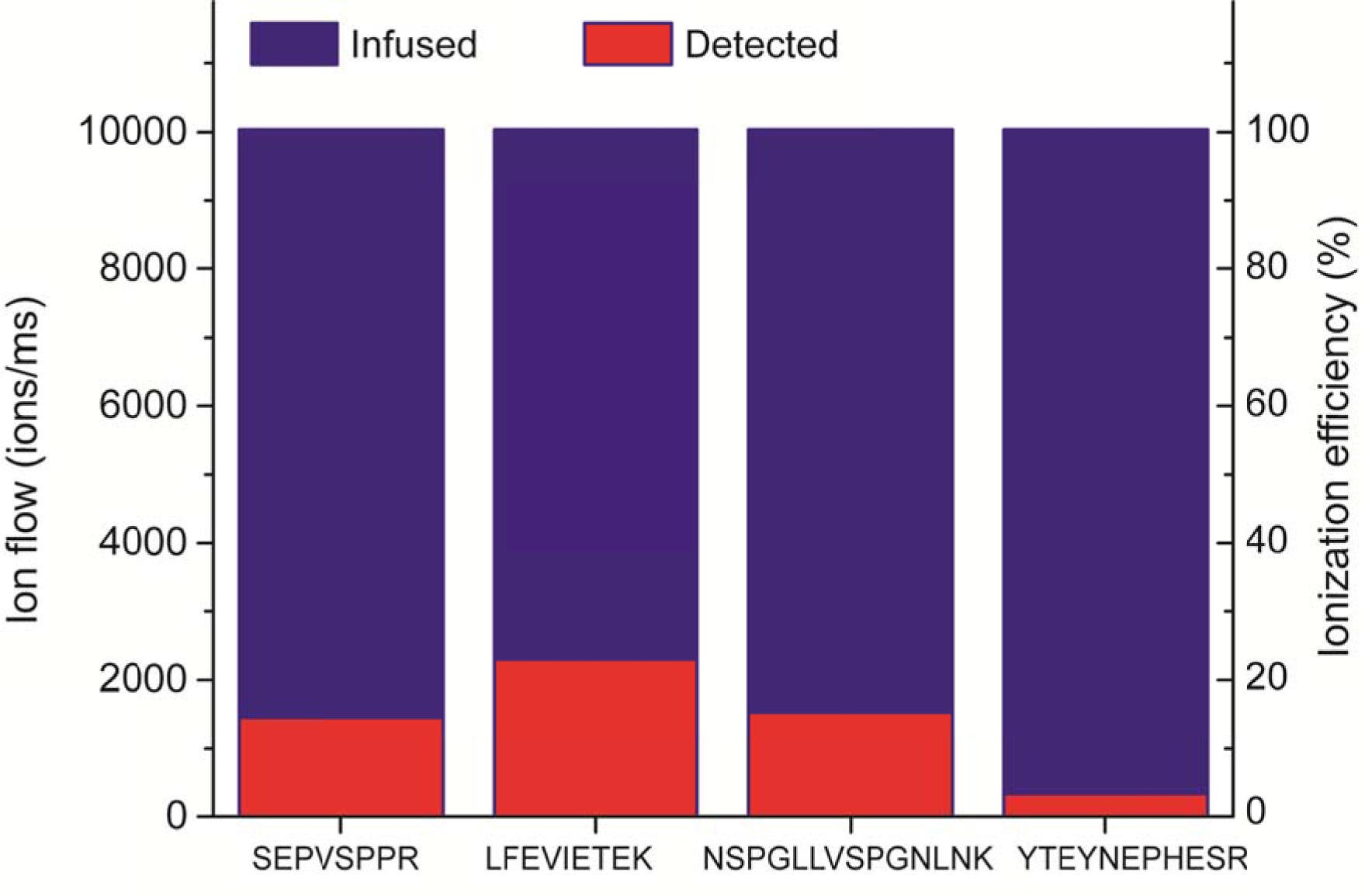
Ionization efficiency estimation for four target peptides, obtained from optimized 2-3 μm tip emitters.

**Supplementary Figure 3.**
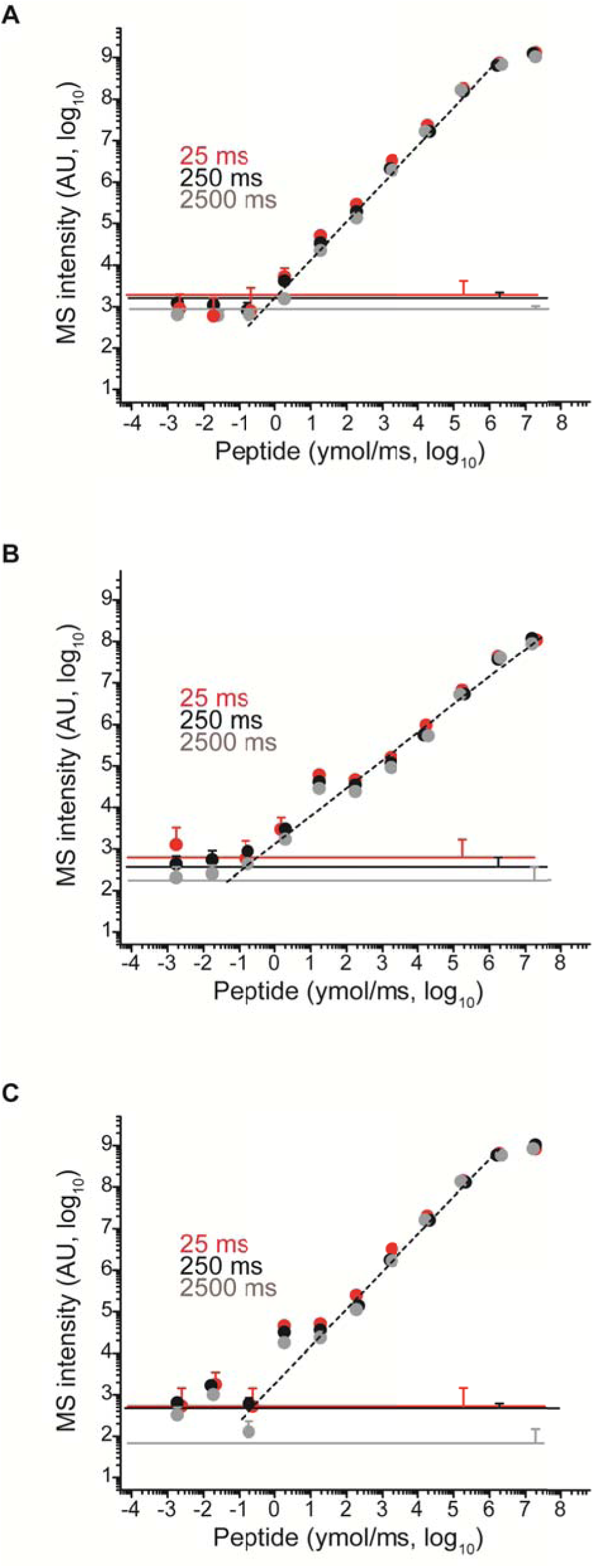
AIM acquisition enables detection of 2-5 yoctomoles/ms of target peptide serially diluted in neat solvent, and seven order of magnitude of linear dynamic range. The MS intensity within 10 ppm from the theoretical m/z was recorded for peptides SPEVSPPR (A), YTEYNEPHESR (B), and LFEVIETEK (C) with maximum injection time (max IT) 25 (red), 250 (black), and 2500 (grey) ms (n=7, error bars = standard deviation). Noise levels recorded at baseline level (i.e., with no target peptide infused) is displayed for each maximum injection times as continuous line.

**Supplementary Figure 4.**
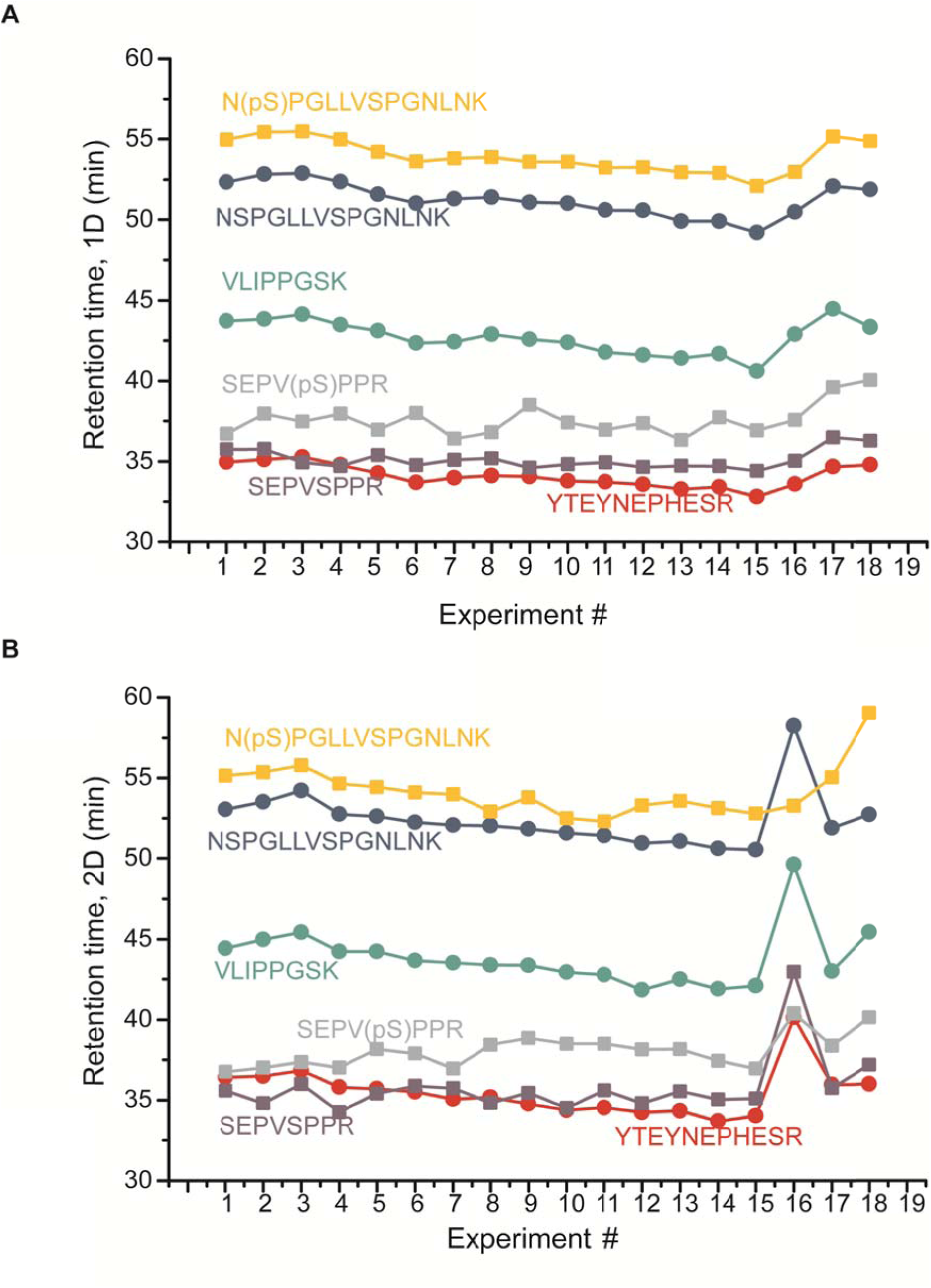
Reverse phase retention times for 6 targets over 18 consecutive sample injections in A) 1D and B) 2D. Outliers runs were not removed to demonstrate true technical variability.

**Supplementary Figure 5.**
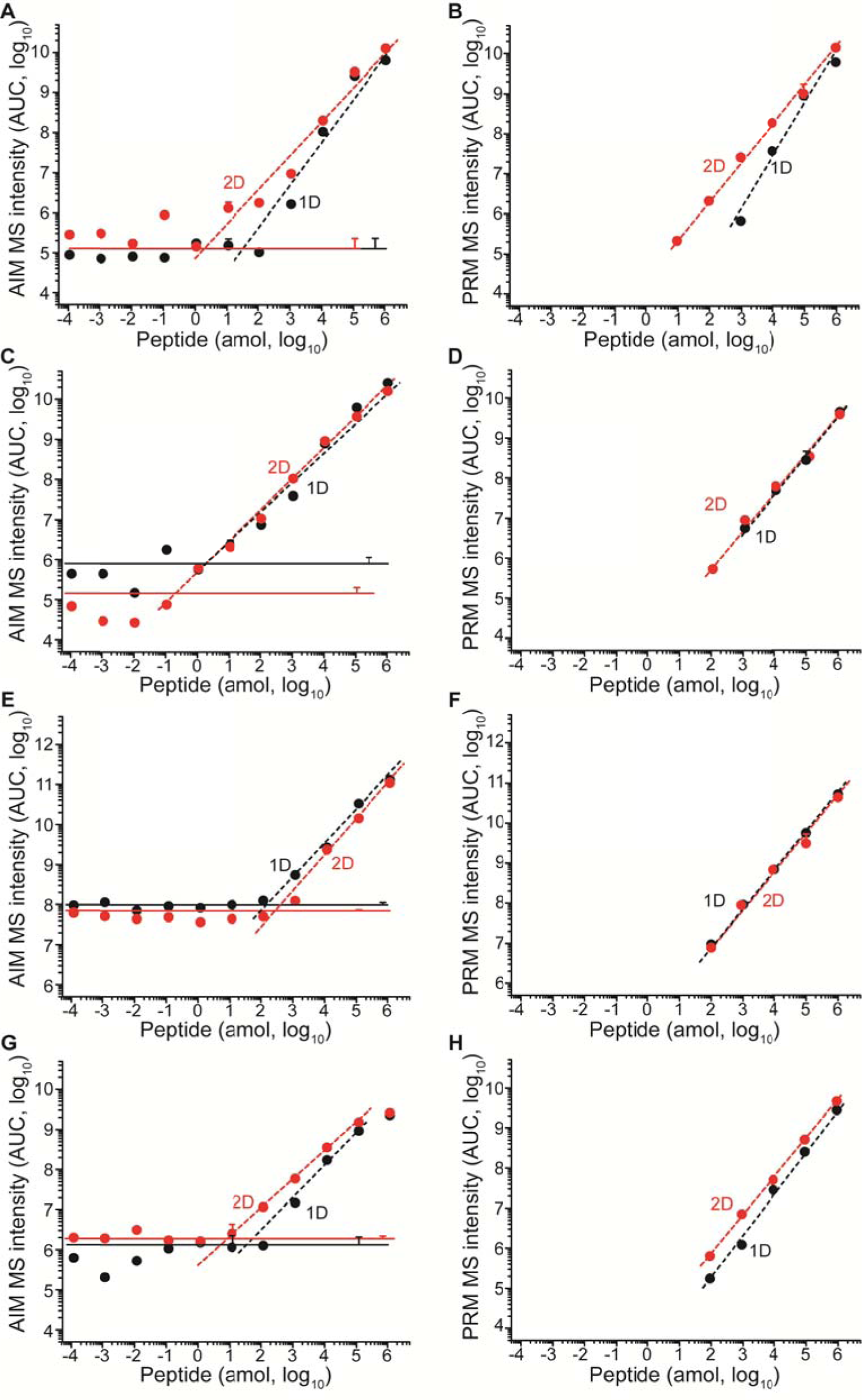
Comparison of sensitivity in AIM and PRM sensitivity following 1D (black) and 2D (red) chromatography. The figures refers to peptides: SEPVSPR (A, B), SEPV(pS)PR (C,D), VLIPGSK (E, F), and YTEYNEPHESR (G, H). Horizontal lines represent average noise levels (n=3) for 1D (black) and 2D (red) in AIM. Error bars represent technical variability as standard deviation (n=3) measured at noise level, 10 amol and 100 fmol. Overlapping data points are horizontally offset when needed for clarity.

**Supplementary Figure 6.**
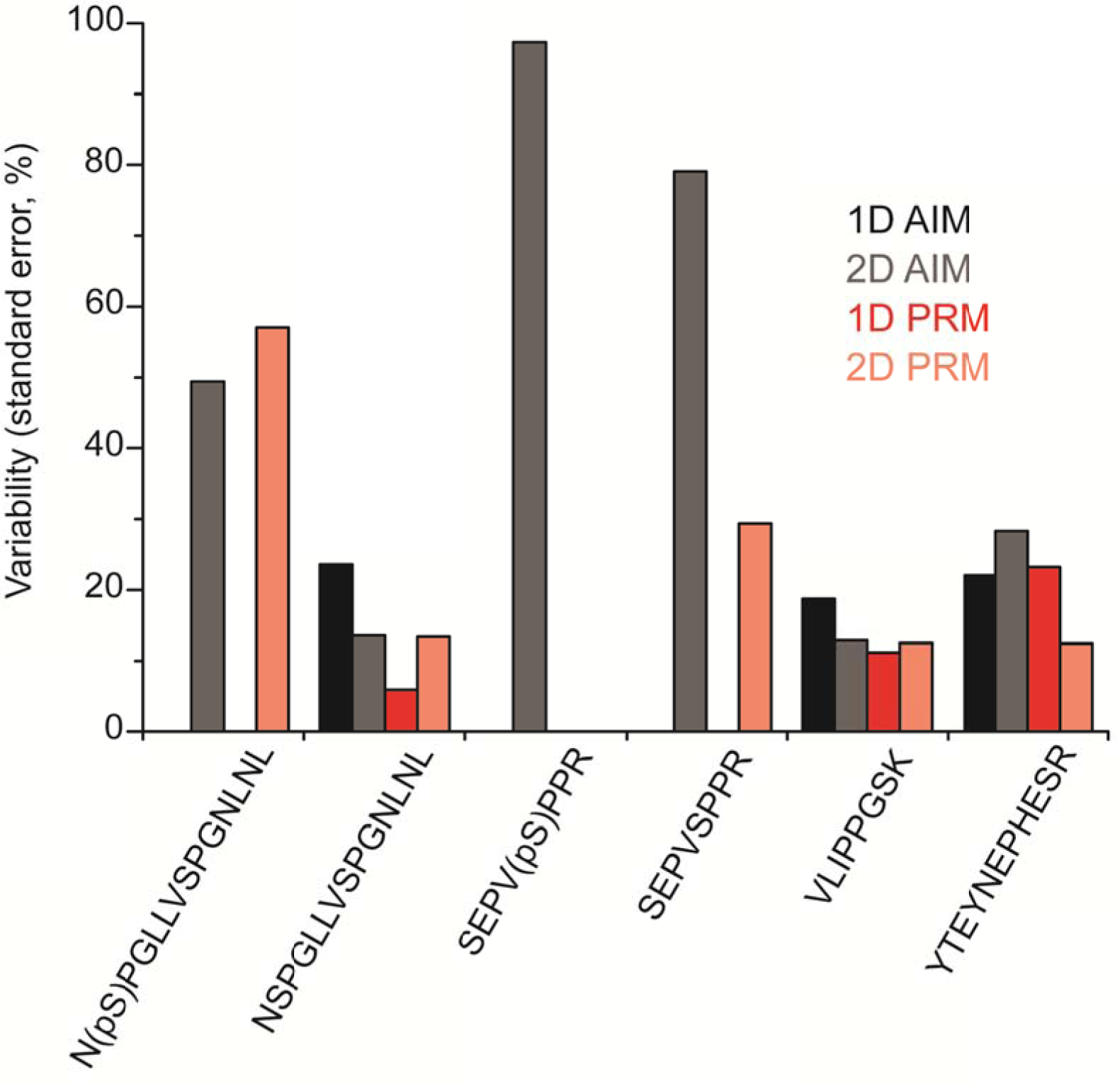
Precision of measurement for AIM and PRM under 1D (shades of grey) and 2D (shades of red), expressed as variability (standard error of the mean) of endogenous peptides quantification.

**Supplementary Figure 7.**
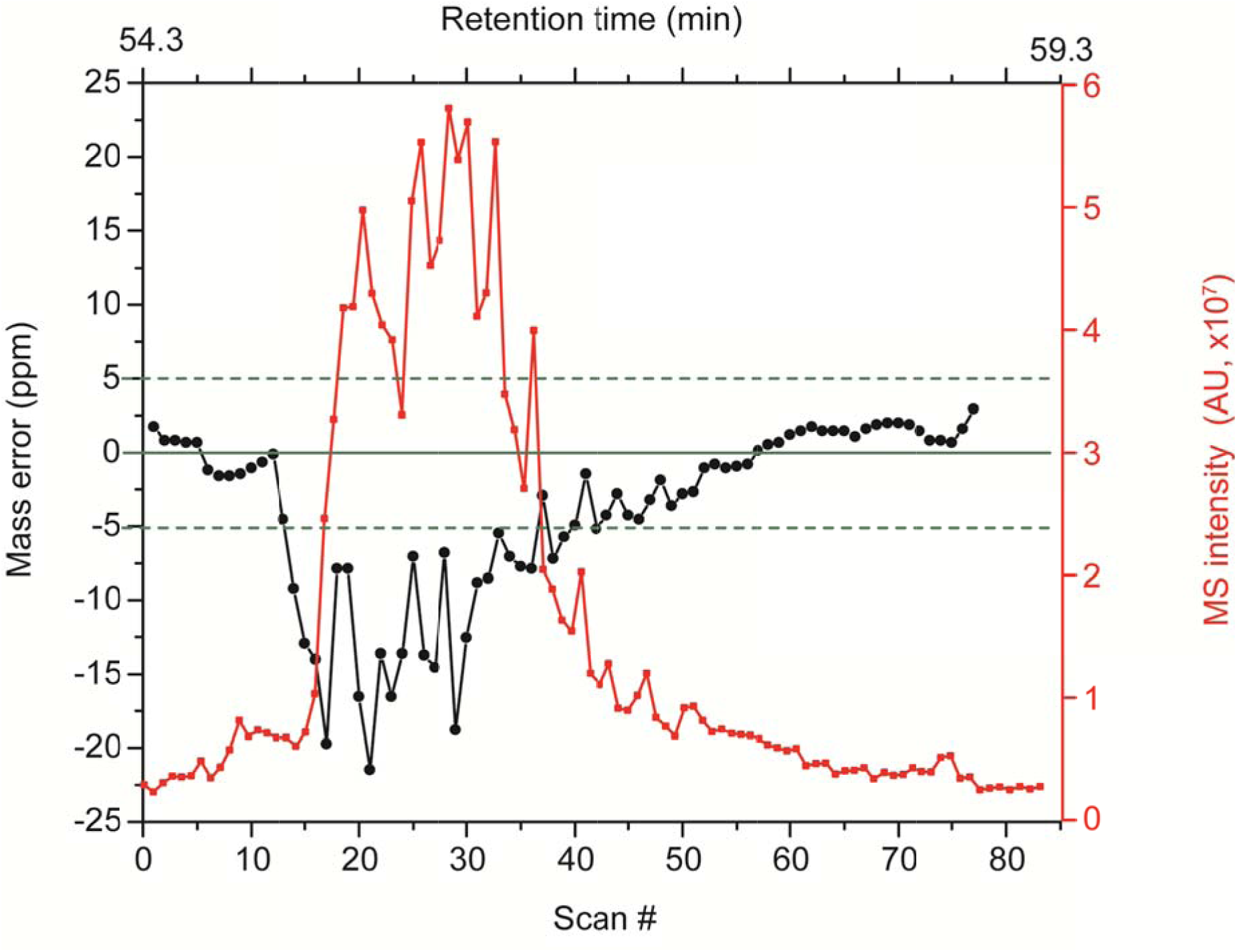
High target flow-rate, here demonstrated with peptide N(pS)PGLLVSPGNLNK at 1 pmol, produced deterioration of mass accuracy (black) and reverse phase resolution (evident as peak broadening to several minutes, red).

## SUPPLEMENTARY MATERIALS

**Supplementary Table 1.** Optical absorbance determination of individual peptide concentration in stock solutions.

**Supplementary Table 2.** MS signal recorded for each target under direct infusion in neat solvent

**Supplementary Table 3.** AIM signal recorded for each target under 1D and 2D.

**Supplementary Table 4.** PRM signal recorded for each target under 1D and 2D.

**Supplementary Methods.** Detailed description and operational parameters of the automated 2D chromatography system.

